# CST does not evict elongating telomerase but prevents initiation by ssDNA binding

**DOI:** 10.1101/2021.08.25.457677

**Authors:** Arthur J. Zaug, Ci Ji Lim, Conner L. Olson, Maria T. Carilli, Karen J. Goodrich, Deborah S. Wuttke, Thomas R. Cech

**Author notes:** To whom correspondence should be addressed.; Correspondence may also be addressed to: Ci Ji Lim., Deborah S. Wuttke. Present address: Ci Ji Lim, Department of Biochemistry, University of Wisconsin Madison, Madison, WI 53706, USA; Maria T. Carilli, Program in Biophysics, Caltech, Pasadena, CA 91125, USA.

## Abstract

The CST complex (CTC1-STN1-TEN1) has been shown to inhibit telomerase extension of the G-strand of telomeres and facilitate the switch to C-strand synthesis by DNA polymerase alpha-primase (pol α-primase). Recently the structure of human CST was solved by cryo-EM, allowing the design of mutant proteins defective in telomeric ssDNA binding and prompting the reexamination of CST inhibition of telomerase. The previous proposal that human CST inhibits telomerase by sequestration of the DNA primer was tested with a series of DNA-binding mutants of CST and modeled by a competitive binding simulation. The DNA-binding mutants had substantially reduced ability to inhibit telomerase, as predicted from their reduced affinity for telomeric DNA. These results provide strong support for the previous primer sequestration model. We then tested whether addition of CST to an ongoing processive telomerase reaction would terminate DNA extension. Pulse-chase telomerase reactions with addition of either wild-type CST or DNA-binding mutants showed that CST has no detectable ability to terminate ongoing telomerase extension in vitro. The same lack of inhibition was observed with or without pol α-primase bound to CST. These results suggest how the switch from telomerase extension to C-strand synthesis may occur.

## INTRODUCTION

Telomerase, a ribonucleoprotein enzyme, comprises a template-containing RNA (1), a reverse transcriptase protein (2), and accessory subunits that differ among ciliates, vertebrates, and yeast (3,4). By maintaining chromosomal telomere length, telomerase allows continuous proliferation of stem cells and cancer cells. The last decades have witnessed substantial progress in understanding telomerase’s enzymatic mechanism, biogenesis, recruitment to telomeres, and three-dimensional structure (5–8). At the same time, research has begun to shed light on the synthesis of the C-rich strand of the telomere (9–14).

Key to the switch from telomeric G-strand synthesis by telomerase to C-strand synthesis by pol α-primase is the CST complex, consisting of CTC1, STN1, and TEN1 (15,16). CST binds single-stranded DNA (ssDNA) with some specificity for the telomeric sequence (15, 17,18). CST prevents telomerase re-initiation by sequestering the 3’ end of the telomeric primer (17). It also directly binds pol α-primase and acts as a cofactor for stimulation of pol α-primase activity (19).

Structures of various domains and subcomplexes of CST were solved by X-ray crystallography in the Skordalakes lab (20,21). The cryo-EM structure then showed how these domains were incorporated into the heterotrimer and revealed a binding site for single-stranded telomeric DNA in the CTC1 subunit (22). Unexpectedly, these heterotrimers can also self-assemble into a 2 megadalton decameric supercomplex with an overall double-ring structure (22). Although data support the existence of the decamer in cells, it remains challenging to ascertain which functions (or additional functions) of CST are accomplished by the heterotrimer versus the decamer (22).

In this work, we utilize human CST DNA-binding mutant proteins that maintain assembly of the heterotrimeric complex but have reduced affinity for telomeric DNA. We find that these mutant proteins have reduced ability to inhibit initiation of telomerase extension. Quantitative profiles of telomerase inhibition as a function of added CST were well fit by an exact treatment of competitive primer binding using experimentally validated binding constants. This analysis provides strong support for the primer sequestration model of Chen, Redon and Lingner (17). We then tested whether CST could terminate ongoing extension of telomeric DNA by telomerase, which would provide a powerful mechanism to switch from G-strand synthesis to C-strand synthesis. However, we show that such termination does not occur to an appreciable extent under multiple conditions in vitro. Together, our data support the model where CST primarily blocks telomerase through primer sequestration, with the switch from telomerase G-strand synthesis to pol α-primase C-strand synthesis occurring either passively or facilitated by factors beyond the telomerase holoenzyme, the CST complex, and pol α-primase.

## MATERIALS AND METHODS

### Reagents

Other than stated, we purchased chemicals from Sigma-Aldrich (St. Louis, MO) or Thermo Fisher Scientific (Waltham, MA), DNA modifying enzymes from New England Biolabs (NEB, Ipswich, MA) and DNA oligonucleotides from Integrated DNA Technologies (IDT, Coralville, IA). The 18-nucleotide 3xTEL DNA is 5⍰-TTAGGGTTAGGGTTAGGG-3⍰. For fluorescence polarization binding assays, 3xTEL was 5⍰ 6-carboxyfluorescein labeled by IDT.

### Biological resources

The pcDNA mammalian expression vector (V79020, Thermo Fisher Scientific) was used to clone cDNAs expressing the human CST subunits. The CTC1 cDNA (MGC: 133331) has a 3xFLAG tag, STN1 cDNA (MGC: 2472) a Myc tag, and TEN1 cDNA (MGC: 54300) a HA tag, all three tags residing on the N-termini of the proteins. CTC1 mutagenesis was performed using standard DNA mutagenesis protocol and confirmed by sequencing the gene. HEK239T cells (CRL-1573, ATCC, Manassas, VA) were cultured in DMEM medium supplemented with 2 mM L-glutamine, 1% penicillin/streptomycin, and 10% fetal bovine serum.

### Expression and purification of proteins in human cultured cells

The three plasmids encoding the CST subunits were transfected into HEK293T cells at 1:1:1 molar ratio using lipofectamine 2000 (11668019, Thermo Fisher Scientific). The cells were further expanded (typically three-fold) for 24 h after transfection and then harvested. The cell pellets were lysed with CHAPS lysis buffer (10 mM Tris-HCl pH 7.5, 1 mM MgCl_2_, 1 mM EGTA, 0.5% CHAPS, 10% glycerol, 5 mM β-mercaptoethanol, 1 mM PMSF) for 45 min at 4 °C on a rotator. The lysate was then clarified by centrifugation at 13,000 x g at 4 °C for 30 min. Anti-FLAG resin (A2220, Sigma-Aldrich) was added to the clarified supernatant and the samples incubated in a rotator for 4 h (or overnight) at 4 °C. The anti-FLAG resins were washed thrice with wash buffer (20 mM HEPES-NaOH pH 8.0, 150 mM NaCl, 2 mM MgCl_2_, 0.2 mM EGTA, 0.1% NP-40, 10 % glycerol, 1 mM TCEP) before elution using wash buffer supplemented with 0.25 mg/mL 3xFLAG peptide (F4799, Sigma-Aldrich). The eluent was then subjected to another round of affinity purification using anti-HA resin (26181, Thermo Fisher Scientific) with similar buffers but 3xFLAG peptide replaced with HA peptide (A6004, APExBIO, Houston, TX) for elution. Purified CST complexes were verified with SDS-PAGE using a silver staining kit (24612, Thermo Fisher Scientific). For experiments that required pol α-primase removal from CST, the NaCl concentration in the wash and elution buffers was raised from 150 mM to 300 mM.

CST protein concentrations were determined by western blot analysis with anti-CTC1 antibody (see next section) using a serial dilution of the HEK-cell CST preparation and a standard curve obtained by serial dilution of an insect cell-purified CST standard. The same quantification with anti-STN1 antibody consistently gave a 3-fold lower protein concentration, possibly because of different post-translational modifications in the HEK cell and insect cell preparations; if the STN1-derived protein concentrations had been used, the *K_d_* and IC_50_ values reported herein would all be 3-fold lower (i.e., tighter binding), but the relative *K_d_* values would be unchanged.

### Western blotting

The presence of CST and pol α-primase subunits in the HEK239T cell-purified CST complexes was analyzed by western blotting. The primary antibodies were anti-FLAG (A8592, Sigma-Aldrich), anti-HA (NB600-362H, Novus Biologicals, Centennial, CO), anti-CTC1 (MABE 1103, EMD Millipore, Burlington, MA), anti-STN1 (NBP2-01006, Novus Biologicals, Centennial, CO), anti-POLA1 (ab31777, Abcam, Cambridge, UK), anti-POLA2 (21778-1-AP, ProteinTech, Rosemont, IL), anti-PRIM1 (10773-1-AP, ProteinTech), and anti-PRIM2 (NBP2-58498, Novus Biologicals). Secondary antibodies used were anti-rabbit (711-035-152, Jackson ImmunoResearch, West Grove, PA) and anti-mouse (715-035-150, Jackson ImmunoResearch). All primary antibodies were diluted 1:1,000 for blotting. The dilution for secondary antibodies was performed according to the manufacturer’s recommendations.

### Electrophoresis mobility shift assay (EMSA)

The 3xTEL oligo was 5⍰ radiolabeled with [γ-^32^P]ATP (NEG035C005MC, PerkinElmer) using a standard T4 polynucleotide kinase labeling protocol (M0201L, NEB). Each binding reaction (10 μL sample volume) contained 500 counts per min (c.p.m.) of radiolabeled 3xTEL in binding buffer (20 mM HEPES-NaOH pH 8.0, 150 mM NaCl, 2 mM MgCl_2_, 0.2 mM EGTA, 0.1 % NP-40, 10% glycerol, 1 mM DTT) with or without CST added. The binding reactions were incubated on ice for 2h before loading onto a 1X TBE, 0.7 % SeaKem® LE Agarose (50004, Lonza Group, Basel, Switzerland) agarose gel. Gel electrophoresis was performed in a cold room (4 °C) for 1.5 h at 6.6 volts/cm. The gels were dried on Hybond N+ (RPN303B, Cytiva Amersham™, Little Chalfont, UK) and 2 pieces of 3MM chromatography paper (3030917, Cytiva Whatman™) at 80°C for 1.25 h. They were then exposed to a phosphorimager screen overnight. The screen was imaged with a Typhoon FLA9500 scanner (GE Lifesciences). The fraction of the DNA bound *θ* by dividing the counts from the gel-shifted band(s) over total counts per lane. The apparent dissociation constant, *K_d,app._*, was then determined from fitting the fraction bound values to the following Hill equation,

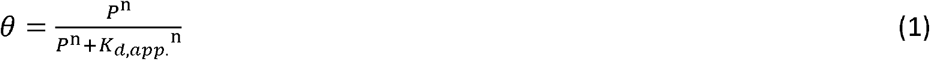

Where P is the CST protein concentration and n is the Hill coefficient.

### Fluorescent polarization (FP) binding assay

Each binding reaction (20 μL sample volume) contained 750 pM of fluorescently labeled 3xTEL oligo in either EMSA binding buffer (for telomerase) or telomerase binding buffer (for CST). Serial dilutions of binding reactions were set up in a 384-well plate (Cat No: 3575, Corning Inc., Corning, NY). Control wells with only binding buffer were also included in each experiment. The binding reactions were incubated for 1.5-2 h at room temperature in the dark. Fluorescent intensity (parallel and perpendicular polarization) of each reaction were measured using a ClarioStar Plus FP plate reader (BMG Labtech, Ortenberg, Germany) and fluorescent anisotropy values of each protein titration were calculated. *K*_d,app._ was determined by fitting the anisotropy value (*F_A_*) to the quadratic equation for single site binding by non-linear least squares fitting,

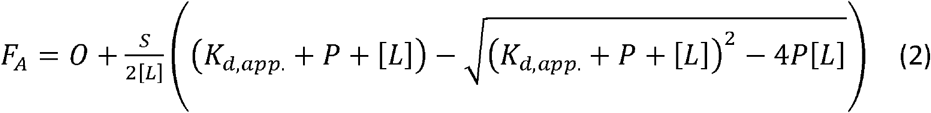

Where *O* is the minimum anisotropy observed, *S* is the difference between the maximum and minimum anisotropy observed, *P* is the concentration of protein, and *[L]* is the concentration of DNA. Averages calculated are the mean values from experiments.

### Direct telomerase assay

Human telomerase expression and purification followed the protocol of Cristofari and Lingner (23). The telomerase extension assay was performed in 50⍰mM Tris-HCl pH 8.0, 50⍰mM KCl, 75⍰mM NaCl (some brought in with CST and the remainder supplemented), 2⍰mM MgCl_2_, 1⍰mM spermidine, 5⍰mM β-mercaptoethanol, 0.33⍰μM [α-^32^P]dGTP (3000⍰Ci mmol−1), 2.9⍰μM cold dGTP, 0.5⍰mM dATP, and 0.5⍰mM TTP.

For standard experiments, CST, telomerase (2.0 nM), and 3xTEL oligo (10 nM unless indicated otherwise) were incubated at room temperature for 30 min before 3⍰μl of dNTP mix was added to initiate telomerase extension (final reaction volume of 20⍰μl). The samples were incubated at 30⍰°C for 1⍰h (unless indicated otherwise) before adding 100 μl of stop solution (3.6⍰M NH_4_Ac containing 20⍰μg glycogen and 3000⍰c.p.m. of each of three oligonucleotide loading controls, LC1, LC2, and LC3). The samples were ethanol precipitated and then dissolved in 10 μl water plus 10 μl 2x gel loading buffer (0.1× TBE, 93% formamide, 50⍰mM EDTA, 0.05% bromophenol blue, and 0.05% xylene cyanol). 10⍰μl of each sample was loaded on a 10% acrylamide, 7⍰M TBE-urea sequencing gel (pre-run for 45 min at 90 W constant) and electrophoresis was performed at 90 W constant until the bromophenol blue dye was at the bottom of the gel, about 2 h. The gel was then dried and exposed to a storage phosphor screen before imaging.

For experiments in which CST was added to an ongoing telomerase reaction, telomerase and 3xTEL oligo were preincubated at room temperature for 30 min before initiating telomerase extension (by adding dNTP mix). CST proteins were then added to the reaction 2 or 10 min after dNTP addition. For pulse-chase experiments, excess cold dGTP and CST were added to the telomerase reactions immediately after the 10 min time point. Radiolabeled telomerase DNA synthesis products were analyzed by ImageQuant (GE Lifesciences). Telomerase activity was determined by total counts per lane, and processive extension was calculated as counts in high molecular weight products (≥ 10 repeats) divided by total counts per lane. IC_50_ values were determined by fitting the telomerase activity data to the equation

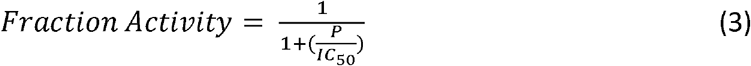

Where *P* is the CST protein concentration.

### Competitive binding modeling and simulation

The following exact mathematical equations for calculating fraction of ligand bound to protein, *θ*, in a competitive binding situation (originally derived by Wang (24) were coded into a python script,

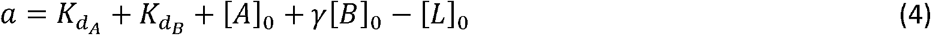

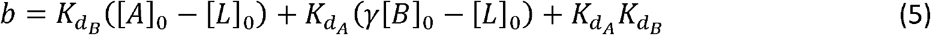

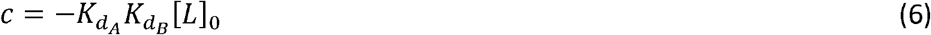

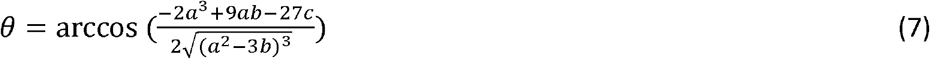

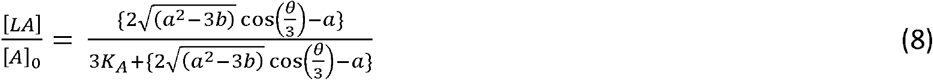

The script was designed to accept user input parameters; 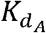 and 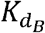, the dissociation constants of the competing binders, (Telomerase and CST, respectively) for the ligand (DNA); [*L*]_0_, the concentration of ligand; [*A*]_0_, the concentration of Telomerase; and [*B*]_0_, a titrated range of initial concentrations of CST.

The final expression calculated is 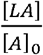, or fraction of telomerase bound to DNA ligand. Normalized fraction bound was then calculated by dividing all values by the value of fraction bound evaluated at [*B*]_0_ nM (in the absence of CST). The equation was also adapted from the original version to accept a manipulatable, unitless γ factor that represented the percent of active CST. This factor was added as a coefficient to concentration of protein B (CST) before calculating normalized fraction bound of ligand to telomerase.

### Fitting of experimental telomerase inhibition data

Best fit curves were generated for experimental competitive binding data. An array of 100 γ values ranging linearly from 0.0 to 2.0 and an array of 100 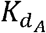 values ranging linearly from 0.0 to 4.0 were created. For every pair of γ and 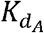 values, a python script was used to calculate the Residual Sum of Squares (RSS) between the exact equation’s predicted fraction bound and the experimentally determined fraction bound under the same conditions according to the following equation,

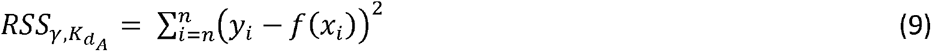

Where *y_i_* is the experimentally determined fraction bound, *f*(*x_i_*) is the exact equation’s prediction of fraction bound under y’s conditions, and *n*, is the total number of experimental data points. The value of 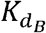 was set at 2.20 nM, the concentration of telomerase at 2.0 nM, and DNA concentrations ranged between 5.0 nM, 10.0 nM, 25.0 nM, 50.0 nM, 100.0 nM, and 200.0 nM, corresponding to telomerase-CST inhibition experiments. 10,000 RSS values were calculated with a minimum value of 0.499 and maximum value of 28.1. Error space was visualized with a 2D heat map corresponding to RSS values for each γ, 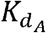 color was set to correspond to the minimum RSS value and the brightest color was set to twice of the saturation RSS value.

The pair with the lowest RSS was then used to generate best fit curves and plotted with experimental data. Best fit curves and heat map were generated using the python Matplotlib graphics package (25).

## RESULTS

### CST mutant proteins defective in binding ssDNA

The cryo-EM structure of human CST revealed a binding site for four nucleotides (TAGG) of the TTAGGG telomeric repeat in Oligonucleotide/oligosaccharide Binding Folds (OB folds) F and G of the CTC1 subunit (22). Mutagenesis was performed on groups of amino acids (designated “g”), designed to give a substantial reduction in DNA affinity (Figure 1A). A negative control mutant g4.1 switched the charge of two amino acids that are not directly involved in DNA binding. While qualitative DNA binding experiments with some of these mutants have been reported (22), the present studies required quantitative measurements.

**Figure 1.**
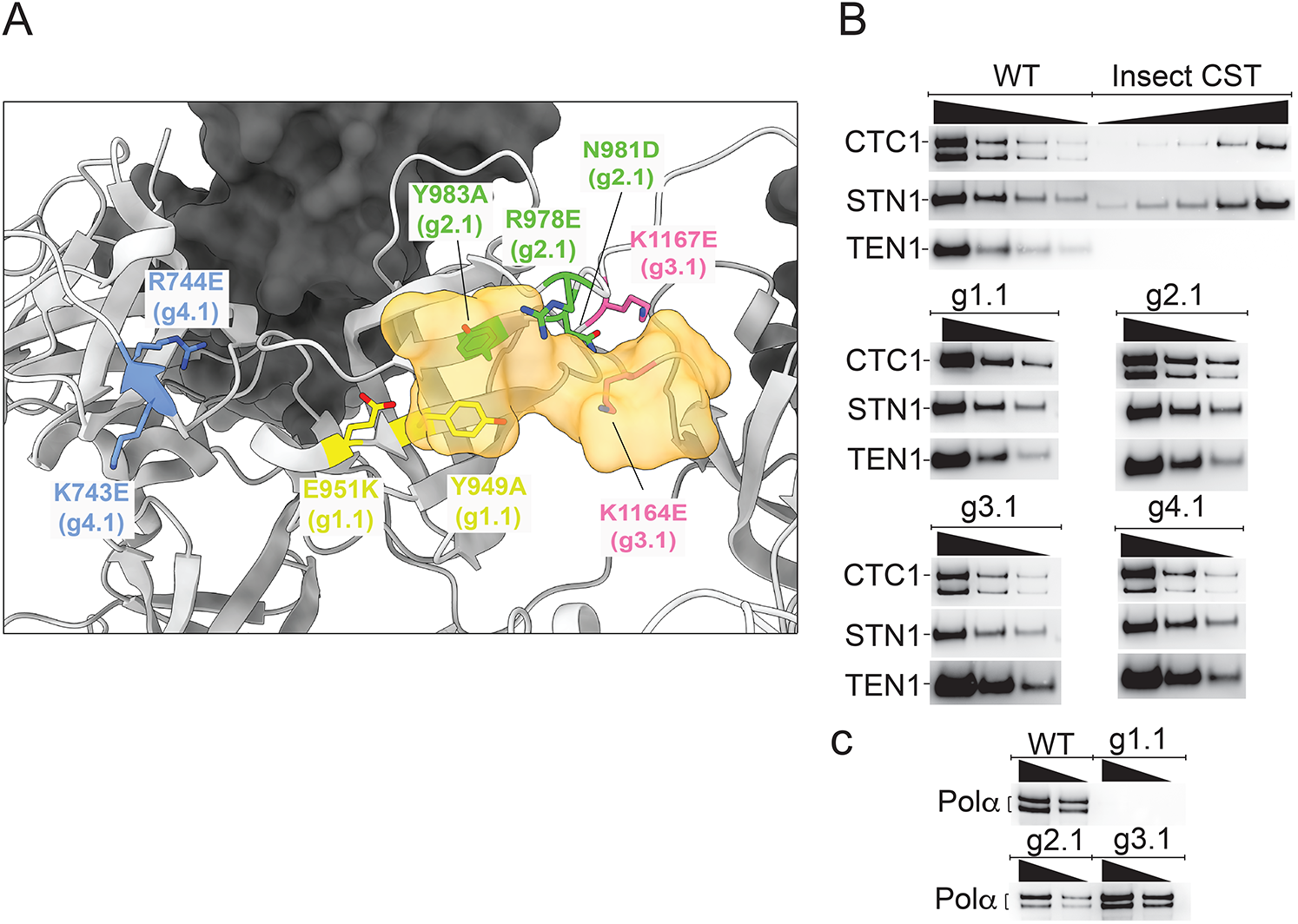
CST DNA-binding mutants maintain subunit assembly and mostly maintain pol α-primase binding. **(A)** Location of mutated amino acids relative to the DNA (half opaque surface representation, orange) in the cryo-EM structure of CST (22). Grey ribbon, CTC1. Dark surface, STN1. **(B)** All mutants maintain assembly of CTC1, STN1 and TEN1 subunits. Insect cell recombinant CST (shown here on WT CST gel) was always included to allow plotting a standard curve to calculate concentration of HEK cell CST. Lack of reliable anti-TEN1 antibody led us to probe the HA-tagged TEN1 with anti-HA antibody; the insect cell TEN1 lacked this tag, so was not revealed. The lower band of CTC1, of unknown origin, was consistently missing in the g1.1 mutant. (C) WT CST and all mutants except g1.1 co-purify with pol α-primase, shown here for the two pol α subunits and in Supplementary Figure S1 for the primase subunits. In panels (B) and (C), wedges indicate successive two-fold dilutions of protein.

The three CST subunits were coexpressed in HEK-293T cells. A double affinity pull-down method relying on a 3xFLAG tag on CTC1 and a HA tag on TEN1 resulted in substantially pure CST complexes (Supplementary Figure S1). The WT and all mutant proteins all assembled stable heterotrimers, as judged by co-IP of the three subunits (Figure 1B). It initially appeared that the protein preparation contained four contaminating polypeptides (Supplementary Figure S1), but mass spectrometry and western blots showed that these were in fact the subunits of pol α-primase, known binding partners of CST (26–29) (Figure 1C). Because pol α-primase was not overexpressed, these subunits are endogenous.

The CTC1 subunit consistently ran as two bands, both containing the N-terminal 3xFLAG tag and the epitope for the CTC1 antibody. The upper band has a molecular weight consistent with full-length CTC1 (135 kDa), while the lower band X (ca. 114 kDa) is of unknown origin. Interestingly, mutant g1.1 was bereft of pol α-primase and of the faster-migrating CTC1 species, providing a useful tool to test whether these components affect DNA binding. Pol α-primase binding appears to be required for nuclear localization of CST (29), but how this could be related to the absence of the smaller CST isoform is unclear.

DNA binding affinity was assessed by both Electrophoretic Mobility Shift Assays (EMSA) and Fluorescence Polarization (FP). Each technique has its advantages, the EMSA allowing detection of a single or multiple bound species, and the FP being more of a true equilibrium technique. For practical reasons, the EMSA was done at 4 °C and the FP at 22 °C, so one would not expect the apparent dissociation constant (*K_d,app_*) values to be the same, but the trends seen with the mutants should be consistent between assays.

Sample EMSA data are shown in Figure 2A. The three mutants designed to be defective in DNA binding bound the 3xTEL DNA probe at much higher protein concentrations than the WT or g4.1 control mutant. Furthermore, the DNA binding-defective mutants all showed at least two DNA-bound complexes on the native agarose gel. The species with the greater retardation had an electrophoretic mobility similar to that of the bound species seen with WT CST and the g4.1 mutant, while the new species ran at an intermediate mobility. It seems unlikely that the intermediate species contains a subcomplex rather than a complete CST heterotrimer, because the three subunits remained associated during immunopurification (Figure 1B) and subcomplexes do not have such high EMSA mobility (30) or structural stability (30,31). In any case, the DNA-binding mutants displayed a 30-50 fold reduction in affinity to the 3xTEL ssDNA, while the negative control had a *K_d,app_* similar to that of WT CST (Figure 2B and Table 1). The curve fits gave Hill coefficients of 1.02 n = 9 experiments) for WT CST and 1.03 0.19 (n = 14 experiments) for the DNA binding mutants, indicating that binding was not cooperative.

**Figure 2.**
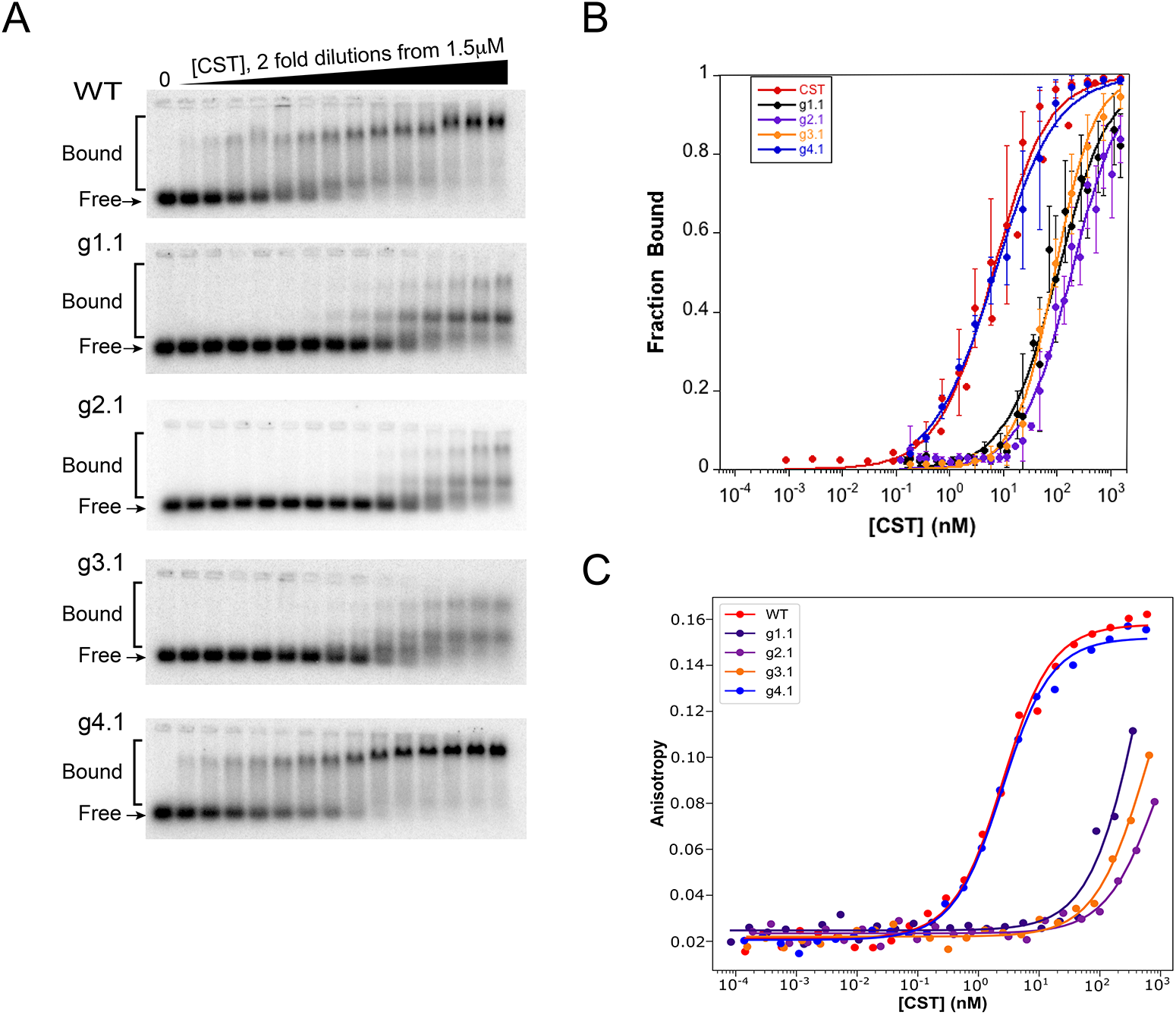
CST mutants show large loss of affinity for telomeric ssDNA. **(A)** Representative EMSA gels of 3xTEL DNA binding by WT and mutant CST proteins. **(B)** Quantification of fraction of DNA bound with error bars representing SD from multiple experiments (see Table 1). **(C)** Representative binding curves from FP assays of 3xTEL binding by WT and mutant CST proteins. Equations 1 and 2 were used for curve fitting.

FP data are plotted in Figure 2C and compiled in Table 1. Consistent with the EMSA data, the DNA-binding mutants showed a large increase in *K_d,app_*, and the g4.1 negative control had a *K_d,app_* similar to that of WT CST. Interestingly, the reduction in affinity for the DNA-binding mutants observed in the FP assays was greater than in the EMSA experiments, 190-360 fold for FP compared to 15-32 fold for EMSA (Table 1). The differences could be explained by the inherent differences of the two assays, with the FP assays being performed at a higher temperature as mentioned above and the FP assays being run at a lower salt concentration to match the telomerase inhibition experiments. Furthermore, the FP assay is better suited than EMSA for measuring binding with weaker binding proteins due to it being a true equilibrium experiment. Overall, though, the trends between the two experiments are consistent.

### CST inhibition of telomerase initiation depends on DNA binding

To compare the ability of various CST complexes to inhibit the initiation of telomerase extension, direct telomerase assays were performed. When telomerase was incubated with the 3xTEL telomeric DNA primer and dNTPs, the 6-nt ladder of extension products characteristic of telomerase was observed, and incorporation of radioactive [α-^32^P]dGTP nucleotides was linear for at least two hours (Figure 3A,B). When WT CST was preincubated for 30 min with telomerase and the primer, the pattern of extension products was unchanged but the intensity of the bands decreased (Figure 3C). The decrease depended on the concentration of CST, with an IC_50_ = 62 5 nM (range of two experiments, 10 nM DNA primer) (Figure 3D). As expected, the IC_50_ increased with increasing DNA primer concentration (Figure 3E, Supplementary Figure S2).

**Figure 3.**
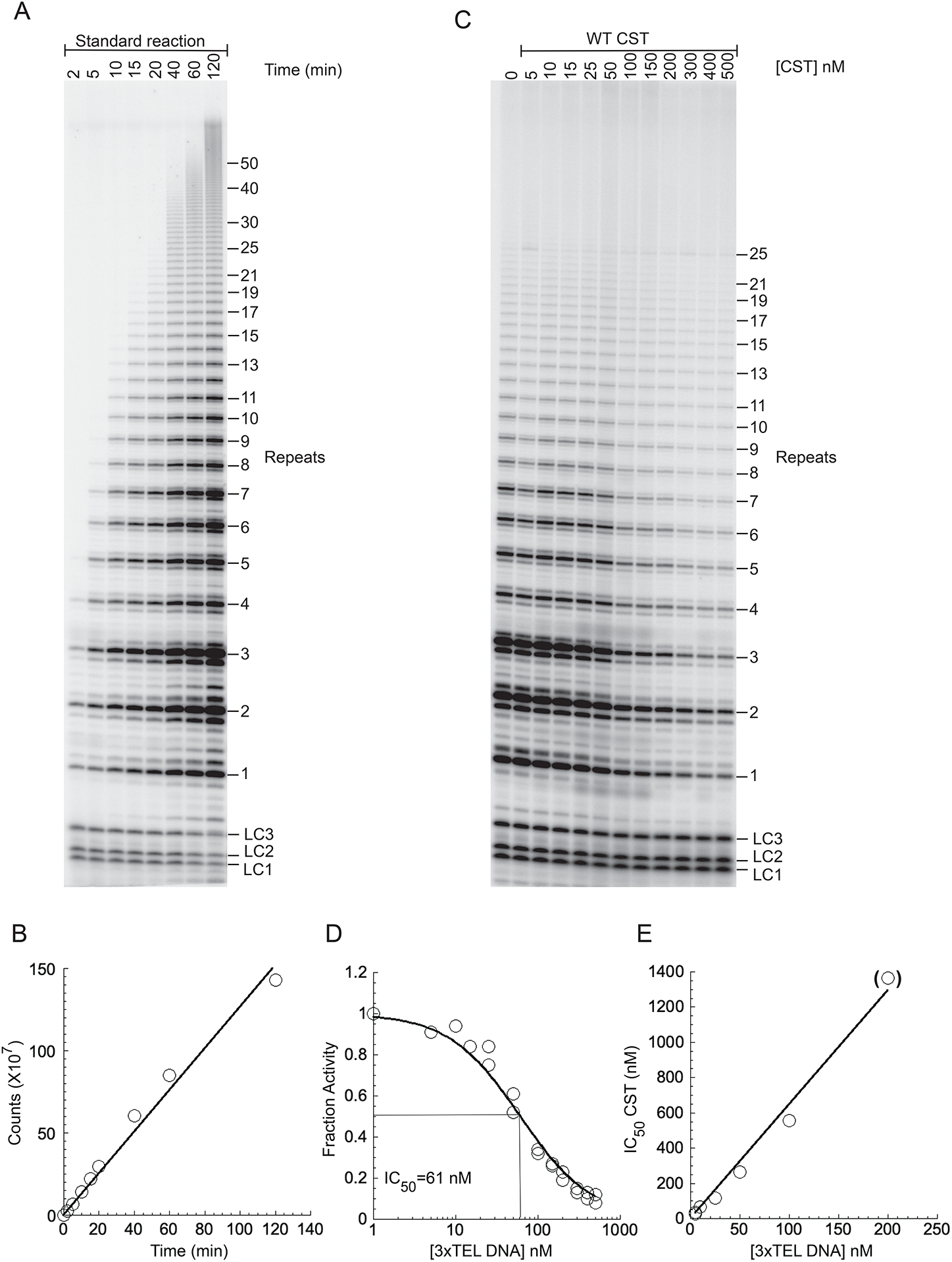
CST inhibits telomerase initiation. **(A)** Direct assay with telomerase immunopurified from HeLa cells, unlabeled 10 nM 3xTEL DNA primer, and nucleotides including [^32^P]dGTP. LC1, LC2 and LC3 are oligonucleotides added as loading controls. **(B)** Radioactivity incorporated into telomerase products as a function of reaction time. **(C)** Telomerase assay for 1 h with the DNA primer pre-incubated with WT CST at the concentrations shown. (D) Counts incorporated into telomerase products normalized to the counts incorporated in the absence of CST. Data from two technical replicates included and fit to equation 3. **(E)** Dependence of IC_50_ on the concentration of 3xTEL DNA in the reaction. Parentheses on last data point indicate that it is underdetermined, because only partial inhibition was achieved at the highest CST concentration.

When the g1.1, g2.1, and g3.1 DNA-binding mutants of CST were added to the telomerase reaction, the inhibition required much higher CST concentrations (Supplementary Figure S3). In more extensive studies of the g2.1 and g3.1 mutants, weak inhibition was observed at low primer concentrations (IC_50_ ~1000 nM), but with 100 nM primer, no inhibition was observed even at 1000 nM CST (Supplementary Figures S4 and S5). Thus, inhibition of the initiation of telomerase activity by CST is dependent on CST’s DNA-binding activity. The negative control g4.1 mutant showed robust inhibition, with an IC_50_ = 10 nM at 10 nM DNA primer and IC_50_ = 471 nM at 100 nM DNA primer (Supplementary Figure S6), similar to WT CST.

Because these inhibition reactions involve two tight-binding entities (telomerase and CST) competing for binding to the DNA primer, it is not immediately apparent how IC_50_ relates to *K_d_*. Thus, in the next section we utilize mathematical modeling to simulate the inhibition curves and fit them to our experimental data.

### Simulating telomerase inhibition by CST with the exact competitive binding expression

The DNA primer sequestration model for telomerase inhibition can be considered a competitive binding equilibrium between CST and telomerase for the DNA primer (17). The schema in Figure 4A illustrates how this competition is governed by dissociation constants defined independently for CST and telomerase based on the free and bound concentrations of all species.

**Figure 4.**
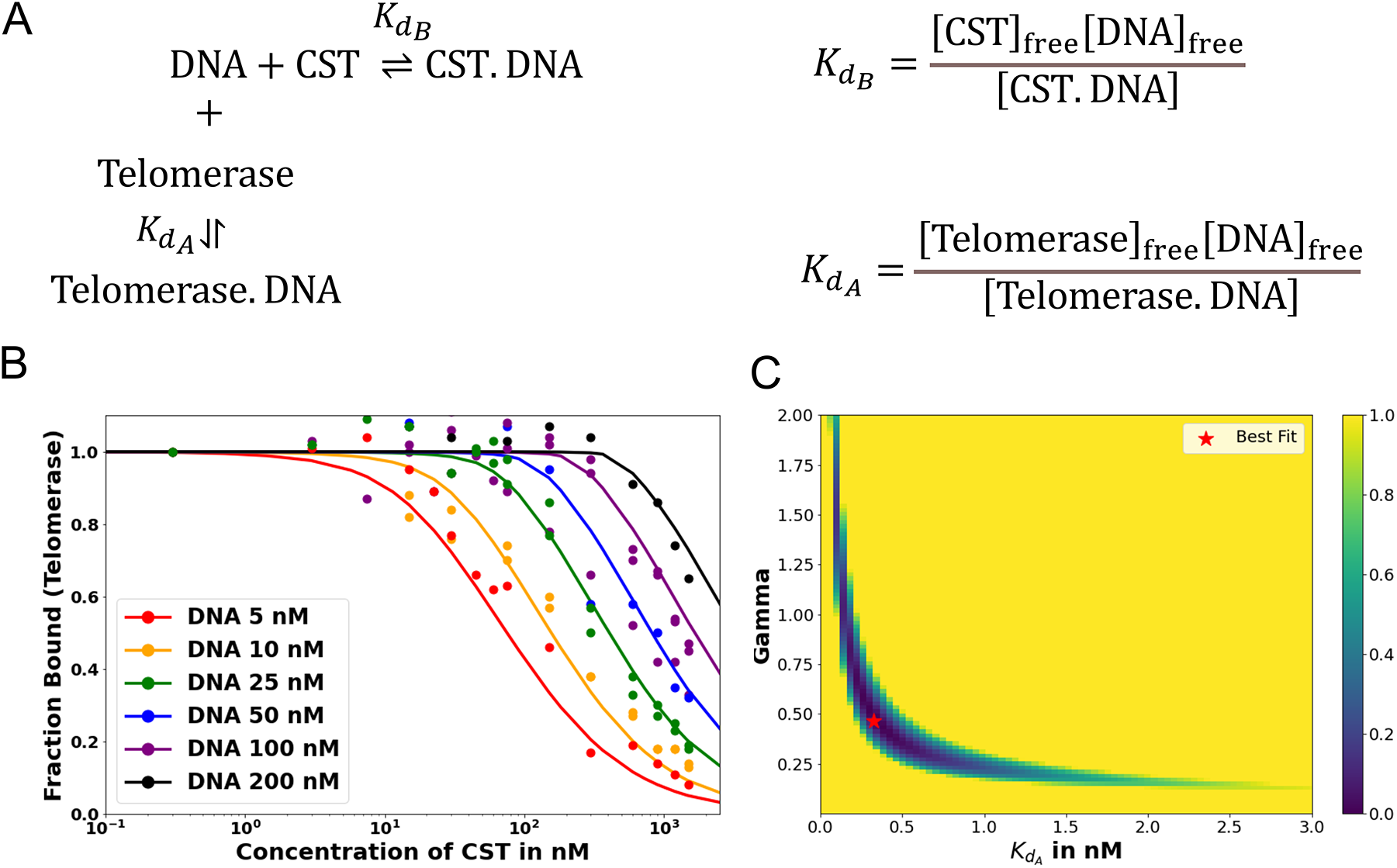
Fitting telomerase inhibition data. **(A)** Schema of competitive binding mechanism and relevant equilibrium equations. **(B)** Best fit telomerase inhibition curves. Telomerase inhibition assay data (points) plotted with predicted fraction bound curves for experimental DNA concentrations generated with optimized *K_d,A_* and γ values of 0.324 nM and 0.465, respectively. Inhibition by WT CST. **(C)** Heat map of error values. Error values (Residual Sum of Squares, see Methods) as a function of γ and *K_d_* Telomerase-DNA pairs with pair of best fit indicated. Value of error colored per bar on right.

The concentrations of DNA, CST, and telomerase relative to the magnitudes of *K_d_*s of CST and telomerase for DNA in our telomerase activity assay were such that the common simplifying assumptions to the competitive binding scenario were not applicable. We thus used the exact, assumption-free mathematical expression for competitive binding situations derived by Wang (24). This exact expression was coded into a python script to accept a series of manipulatable parameters: dissociation constants for two competing binders (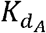 and 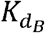), concentration of ligand, initial concentration of constant binder A (telomerase), and a range of initial concentrations of titrated protein B (CST). The script calculated normalized fraction bound of telomerase to ligand at each input concentration of titrated CST (see Materials and Methods). The fraction bound values were then plotted against CST concentration on a logarithmic scale.

A series of test simulations were performed to verify that this exact expression behaved as expected (Supplementary Figure S7). With five different ligand concentrations (5, 25, 50, 100 and 200 nM) and a 2 nM concentration of telomerase (conditions of the telomerase activity assay), we simulated the behavior of the exact expression at several ratios of dissociation constant of CST 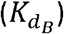 to dissociation constant of telomerase 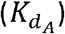. The concentration of titrated CST ranged from 0 to 1,000 nM. As expected, the higher the ratio of CST-DNA 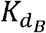 to telomerase-DNA 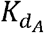 (the weaker CST’s affinity for DNA relative to telomerase’s affinity for DNA), the more the normalized fraction bound telomerase curves shifted to the right (more CST was required to sequester DNA from telomerase). This is apparent in Supplementary Figure S7 by comparing the family of curves in each panel.

Also as expected, increasing the initial concentration of DNA shifted the fraction bound curve to the right regardless of the ratio of CST-DNA 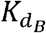 to telomerase-DNA 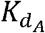 (compare the different curves within each panel in Supplementary Figure S7). With more DNA in solution, more CST was necessary to bind all available DNA, thereby sequestering it from telomerase. These simulations confirmed that exact expression behaved as expected. We then proceeded to fit the expression to the telomerase inhibition data to determine if the inhibition of telomerase action by CST could be successfully modelled as a competitive binding situation.

### Fitting experimental telomerase inhibition data to the exact competitive binding expression

The exact competitive binding expression was then used to fit the telomerase inhibition data collected at a range of DNA concentrations to determine if the primer sequestration model accurately described telomerase inhibition by CST. Several input conditions for the expression were known in the telomerase inhibition activity assays or determined experimentally through independent methods, including initial DNA concentration, telomerase concentration, titrated CST concentrations, and CST-DNA 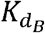 (determined to be 2.2 ± 0.28 nM via FP, Figure 2C). This left two unknown variables to be fit using our model, the telomerase-DNA 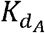 and the active concentration of CST. For the percent of active CST, we modified the exact expression derived by Wang (24) to include a manipulatable, unitless gamma factor as a coefficient on the concentration of CST that represented the percent of active CST.

To simultaneously fit all the experimental data, we tested a set of 10,000 Telomerase-DNA K_DB_s and gamma pairs (with limits set by reasonable physical approximations) to find the pair that minimized error between fraction bound values predicted by the exact expression and values determined in the telomerase inhibition assays. The error between predicted and experimental fraction bound was quantified by calculating the Residual Sum of Squares (RSS) for each gamma, 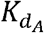 pair (see Materials and Methods).

Using this strategy, optimized values were found to be a 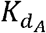 of 0.324 nM and a gamma of 0.465. The optimized fit 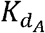 for telomerase-DNA binding was similar to that of 0.54 + 0.25 nM independently determined for our telomerase enzyme by FP (Table 1 and Supplementary Figure S8). These values were then used to generate the best fit curves plotted with telomerase inhibition data (Figure 4B). Note that even at the lowest [DNA] = 5 nM, the calculated IC_50_ is ~70 nM CST, far greater than the binding constant 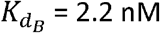. The high IC_50_ concentration matches both experimental and fit telomerase 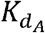 values, which show telomerase has roughly a 4-7 fold higher affinity to DNA than CST.

While a 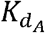 of 0.324 nM and a gamma of 0.465 resulted in lowest error, other pairs had similarly low error values: the RSS values for each pair are represented in a heat map (Figure 4C). This spread of low error values indicated that there is a range of pairs that similarly fit the inhibition data well. These fits show that our data convincingly support competitive primer binding as a model for telomerase-CST inhibition.

### CST does not inhibit ongoing extension of telomeric DNA by telomerase

As shown above, CST can compete with telomerase for binding the DNA primer and thereby inhibit initiation of G-strand synthesis. A much more powerful mode of CST inhibition might occur if it could attack and disrupt ongoing telomerase extension. However, our data suggested that CST might not affect ongoing extension. First, when WT CST was added to the telomerase reaction at 2 min or at 10 min, the incorporation of radioactivity into telomerase reaction products was largely but not entirely curtailed (Figure 5A,B). Second, existing extension products continued to elongate (Figure 5C). Both observations were consistent with CST inhibiting initiation of new telomerase reactions but not affecting ongoing extension.

**Figure 5.**
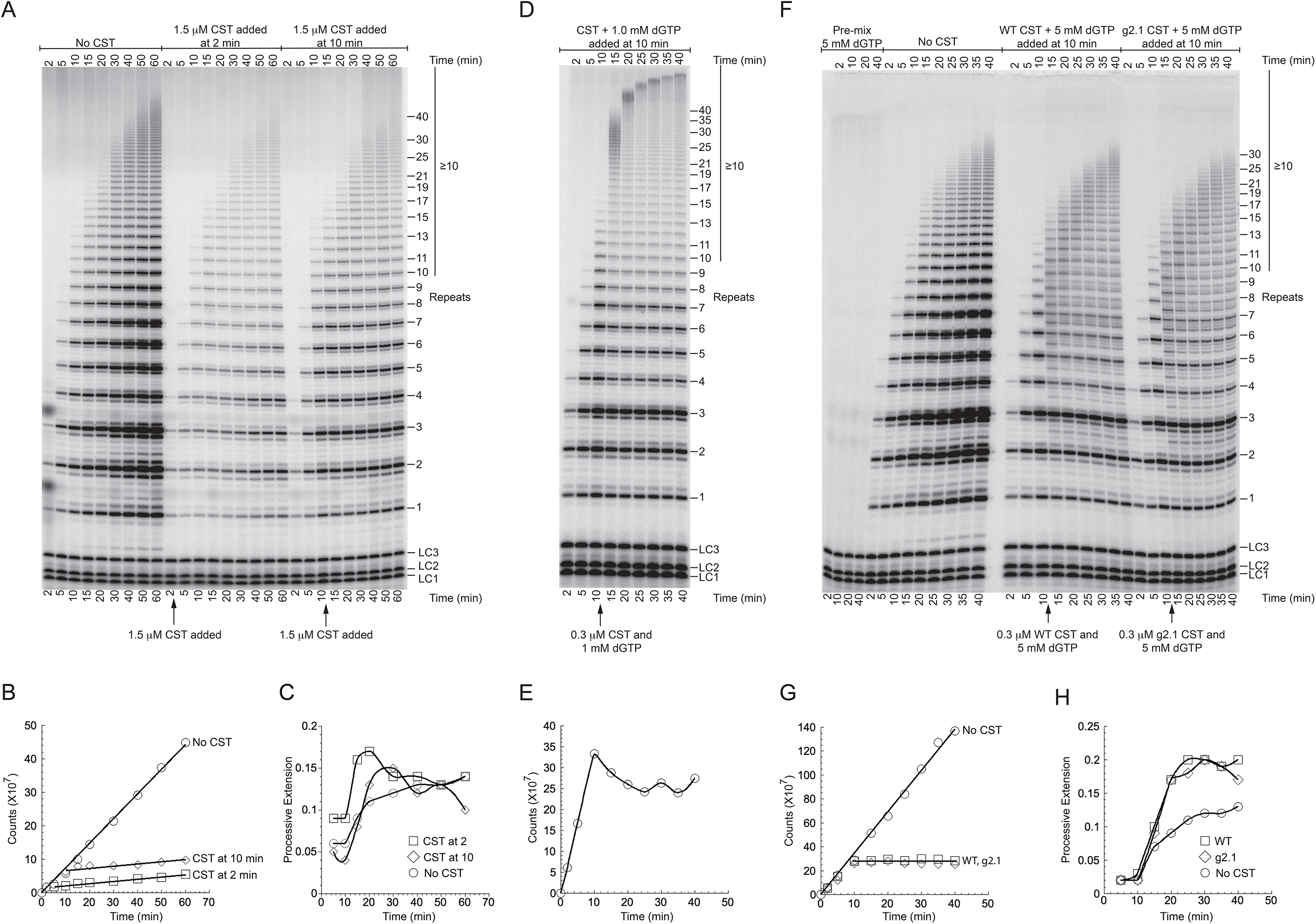
Pulse-chase experiments show that CST does not disrupt ongoing telomerase extension. **(A)** Direct telomerase assay with WT CST added after 2 or 10 min of reaction. All three time courses had the same [^32^P]GTP, so intensity differences are due to inhibition by added CST. **(B)** Radioactivity incorporated into telomerase products as a function of reaction time. **(C)** Long extension products (≥10 repeats, see bar to right of panel a) as a fraction of total incorporation. Data points are connected to aid visualization. **(D)** Pulse-chase experiment in which excess unlabeled dGTP was added along with CST to restrict observed products to previously initiated chains. High [dGTP] drives high processivity. **(E)** Radioactivity incorporated into telomerase reaction products stops increasing when unlabeled dGTP is added at 10 min. Data points are connected to aid visualization. **(F)** WT CST and DNA-binding mutant g2.1 CST are equally unable to disrupt ongoing telomerase extension under conditions that cause telomerase stuttering. Pulse-chase experiments with 5 mM unlabeled dGTP instead of the standard 1 mM dGTP. The pre-mix control (first four lanes) shows that the addition of the unlabeled dGTP was sufficient to prevent further labeling of products. **(G)** Radioactivity incorporated into telomerase products in the experiment of panel (F). Symbols for WT and g2.1 CST overlap. **(H)** Long extension products as a fraction of total incorporation in the experiment of panel (F). Data points are connected to aid visualization. The values with no CST addition are lower because small products continue to be initiated, increasing the total counts relative to the long extension products.

The ladder of extension products in telomerase reactions is generally thought to result from processive extension, because the excess of unextended primer in the reactions should act as an internal “chase” and prevent rebinding of telomerase to previously extended products (32,33). To test if the long extension products that continued to accumulate after CST addition were in fact due to processive elongation of previously initiated chains, we performed pulse-chase experiments. Instead of chasing with a 3’-end-blocked DNA primer, which would prevent telomerase reinitiation but would also bind to CST, we added an excess of unlabeled dGTP along with the CST immediately after the 10 min timepoint so that all further nucleotide incorporation would be unlabeled. An example of such an experiment is shown in Figure 5D. Three observations confirmed that the pulse-chase experiment was working as expected. First, incorporation of radiolabel stopped immediately upon the addition of cold dGTP, present in 3000x excess over the radiolabeled dGTP (Figure 5E). Second, smaller extension products at 10 min decreased in intensity at subsequent time points, as they were chased into longer extension products (see, for example, 7, 8 and 9 repeats). Third, longer products continued to accumulate after 10 min. (As expected, the high [dGTP] stimulated telomerase processivity (34–36).) In conclusion, the addition of a saturating amount of WT CST clearly does not inhibit ongoing processive telomerase activity.

To control for possible nonspecific effects of adding protein to the reaction, WT CST was compared side-by-side with the g2.1 DNA-binding mutant. As shown in Figure 5F,G,H, the WT and g2.1 mutant additions gave essentially identical results. Neither of them inhibited further extension of previously initiated primers (Figure 5H). The stuttering banding pattern (synthesis stalls at 3 nt as well as 6 nt in each repeat) is caused by the higher ratio of dGTP:Mg^++^ in this experiment (5 mM dGTP:1 mM MgCl_2_). Thus, even under conditions where telomerase extension is suboptimal, the CST has no detectable ability to inhibit ongoing telomerase extension. Pulse-chase reactions with 1 mM dGTP:1 mM MgCl_2_, which do not give the stuttering pattern, again showed no difference between WT and g2.1 CST (Figure 6C).

**Figure 6.**
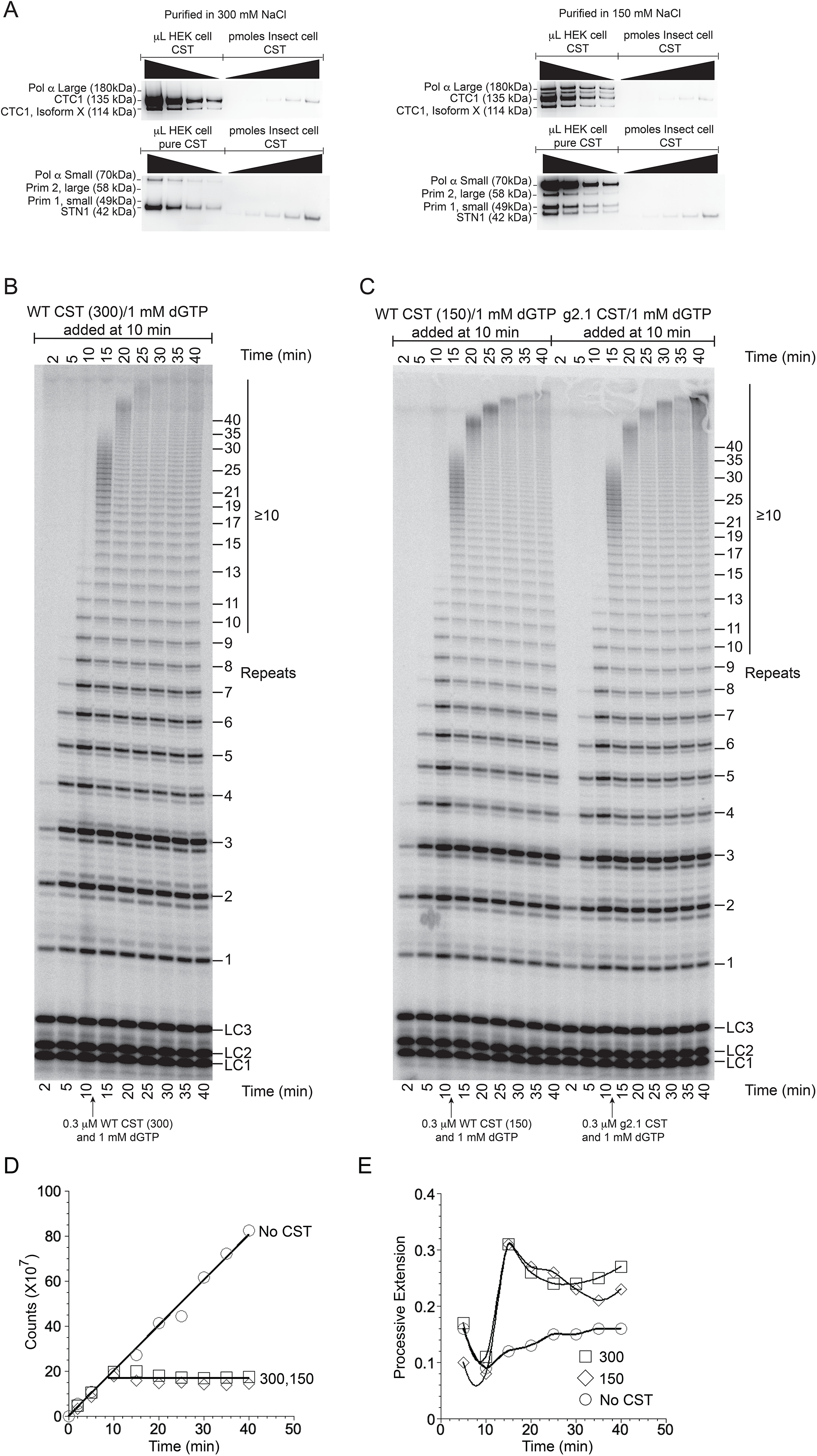
CST inhibits telomerase extension independent of bound pol α-primase. **(A)** Western blot analysis of WT CST purified in standard 150 mM NaCl or in 300 mM NaCl. Top blots probed with antibodies to CTC1 and to the large subunit of pol α. Bottom blots probed with antibodies to STN1, the small subunit of pol α, and the two subunits of primase (Prim 1 and 2). Wedges indicate successive two-fold dilutions of protein. **(B)** Pulse-chase experiment with WT CST purified in 300 mM NaCl. **(C)** Pulse-chase experiments with WT CST purified in 150 mM NaCl (left lanes) and with g2.1 mutant CST purified in 150 mM NaCl (right lanes). **(D)** Radioactivity incorporated into telomerase products as a function of reaction time without chase (No CST) or in the pulse-chase experiments of panel (B) and left half of (C) (300 and 150, respectively). **(E)** Long extension products (≥10 repeats) as a fraction of total incorporation in the same pulse-chase experiments. Data points are connected to aid visualization.

### CST inhibits telomerase extension independent of bound pol α-primase

Pol α-primase copurified with the CST purified from HEK-293T cells. The exception was the g1.1 mutant, which lost pol α-primase association (Figure 1C, Supplementary Figure S1). Because the g1.1 mutant inhibited telomerase activity only at high concentrations, similar to g2.1 and g3.1 which retain pol α-primase, it appeared that pol α-primase was not responsible for the inhibition seen with CST.

We sought an independent test of this hypothesis. We found that immunopurification of the CST under higher salt conditions, 300 mM KCl instead of our standard 150 mM KCl, released the pol α-primase without otherwise affecting the purification of CST (Figure 6A). The 150 mM and 300 mM KCl preparations of CST were compared in a pulse-chase experiment, and they were found to be equivalent: they both prevented further initiation of telomerase, and they both allowed processive extension of pre-initiated chains (Figure 6B-E). Thus, pol α-primase does not appear to be responsible for or to affect CST inhibition of telomerase.

## DISCUSSION

The switch from telomeric G-strand synthesis by telomerase to C-strand synthesis by pol α-primase is a critical step in telomere replication. The CST complex is key in orchestrating this switch, but its mechanism of action is incompletely known. Here we extend the understanding of CST inhibition of telomerase extension in three ways. First, we validate DNA-binding mutants of CST and show that they no longer inhibit telomerase initiation, providing strong additional support for the primer sequestration model. Second, we develop an exact model for CST inhibition of telomerase, which reconciles the *K_d_* values for telomerase and CST binding to the DNA primer with the much higher IC_50_ values obtained from the inhibition curves. Finally, we show that CST does not have the intrinsic ability to evict telomerase from telomeric DNA during a primer-extension reaction.

### An exact model for CST inhibition of telomerase

While CST has previously been suggested to inhibit telomerase through the direct and competitive binding of the telomeric ssDNA ligand, more sophisticated actions by CST could not be ruled out (17). We have determined the extent to which a competitive binding model evaluated with an exact treatment of coupled equilibrium (24) fits the experimental telomerase inhibition profiles. The strength of this approach is that it requires no a *priori* assumptions about relative dissociation constants or limiting values to predict the equilibrium concentrations of all species. This model, however, treats the telomerase-DNA interaction as a straightforward binding interaction, a potentially deleterious oversimplification of the complex enzymatic machinery. Phenomenon such as partial dissociations, multiple binding modes, processivity and change in affinity for an active telomerase are not expected to be accurately encapsulated by a single binding constant. Thus, it is somewhat surprising that this model represents the inhibition data quite well. Furthermore, the fitted value for telomerase-DNA binding affinity (Figure 4B) matches within error to the value obtained independently in a binding experiment (Table 1). This binding affinity (0.52 ± 0.25 nM) is similar to other values reported in the literature measured with alternate methods of 0.5 ± 0.3 nM (37) and 3.3 ± 0.5 nM (38). The high congruence of the model with the data, as well as the consistent telomerase-DNA affinity values obtained, strongly support that the core action of telomerase inhibition by CST is through the mechanism of competitive binding for ligand. This mechanism suggests the possibility that regulation of telomerase action could be achieved by controlling the CST concentration at the telomere.

### How does CST terminate telomerase extension?

Telomerase is unable to extend ssDNA primers in the presence of a saturating amount of CST. This activity can be explained by a simple primer sequestration model (17). That is, CST binding to the primer sequesters it and precludes telomerase binding. Our detailed quantitative analysis with DNA-binding mutants of CST provides strong confirmation of the primer sequestration model with no need to invoke additional CST activities.

How effective might CST inhibition of initiation be for termination of telomerase in vivo? During homeostatic telomere length maintenance, human telomerase is thought to extend most telomeres in each cell cycle, adding about 60 nt to each telomere processively after a single binding event (39). The intrinsic activities of CST determined by in vitro analysis are in complete accord with such a model. Given its ability to sequester the primer and block reinitiation, CST could help restrict telomerase extension to a single round. Because it is unable to evict telomerase from elongating DNA, it would not interfere with the single round of extension.

On the other hand, when telomeres are undergoing net elongation, Zhao et al. (39) report that multiple telomerase molecules act at each telomere; i.e., extension is distributive rather than processive. Perhaps under these conditions there is insufficient CST available at the telomere to prevent telomerase reinitiation.

How does the switch from telomerase synthesis of the telomeric G-strand to pol α-primase synthesis of the C-strand occur? Because we find no evidence that CST can evict telomerase from elongating DNA under multiple in vitro conditions, the switch from G-strand to C-strand synthesis may be passive rather than active. Given its modest processivity, telomerase will terminate spontaneously after adding a limited number of telomeric repeats, measured as ~4 repeats in vitro (40) and ~10 repeats in cells (39). Inhibition occurs when CST then binds the newly extended DNA and prevents telomerase reinitiation. At the same time, CST brings in pol α-primase to initiate C-strand synthesis.

The relative activity of the CST heterotrimeric “monomer” and the 2-megadalton decameric supercomplex in inhibiting telomerase is not addressed by our study. We think that the results presented here pertain to the CST monomer. The HEK cell-based CST purification gives low concentrations of CST, and under these conditions we have not observed the decameric supercomplex (although our evidence suggests that it is present in cells (22)). Given that the decamer is poised to bind ssDNA more aggressively than the monomer (22), we might expect it to have greater telomerase-inhibiting activity than the monomer.

The experimental approaches and computational analysis developed herein provide the groundwork for future studies to test the activity of additional telomere components, such as the shelterin complex, on the switch from telomeric G-strand to C-strand synthesis.

## AVAILABILITY

CST_Inhibits_Telomerase is available in the GitHub repository (https://github.com/mtcarilli/CST_Inhibits_Telomerase).

## SUPPLEMENTARY DATA

Supplementary DATA are available at NAR Online.

## ACKNOWLEDGEMENTS

We thank T. Nahreini for her excellent management of the Biochemistry Cell Culture Facility, CU Boulder. T.R.C. is an investigator of the Howard Hughes Medical Institute.

---

Table 1. Equilibrium dissociation constants for binding of CST and telomerase to 3xTEL DNA by EMSA at 4°C and by FP at room temperature. Each value is the mean SD of *n* independent experiments. Relative *K_d_* values are normalized to WT CST. For the weak-binding CST mutants, the FP binding curves did not go to completion, so the *K_d_* values are approximate.

