## Supplemental Figures for "CST does not evict elongating telomerase but prevents initiation by ssDNA binding"

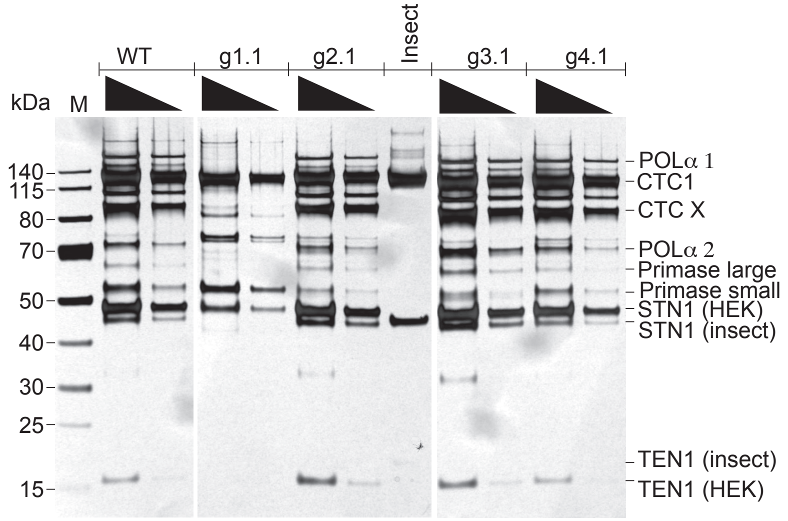


**Supplementary Figure S1**. SDS-PAGE analysis of CST protein preparations. Wedges represent two-fold dilutions. Gels were silver-stained. CTC1 X is the CTC1 isoform described in the text. Western blot analysis confirmed that the band in g1.1 co-migrating with primase small is not primase small. Recombinant CST expressed in insect cells (22) gave STN1 and TEN1 with slightly different mobility than those expressed in HEK293T cells due to different N-terminal tags. TEN1 was close to the limit of detection, so its presence was evaluated by western blotting (Figure 1B)

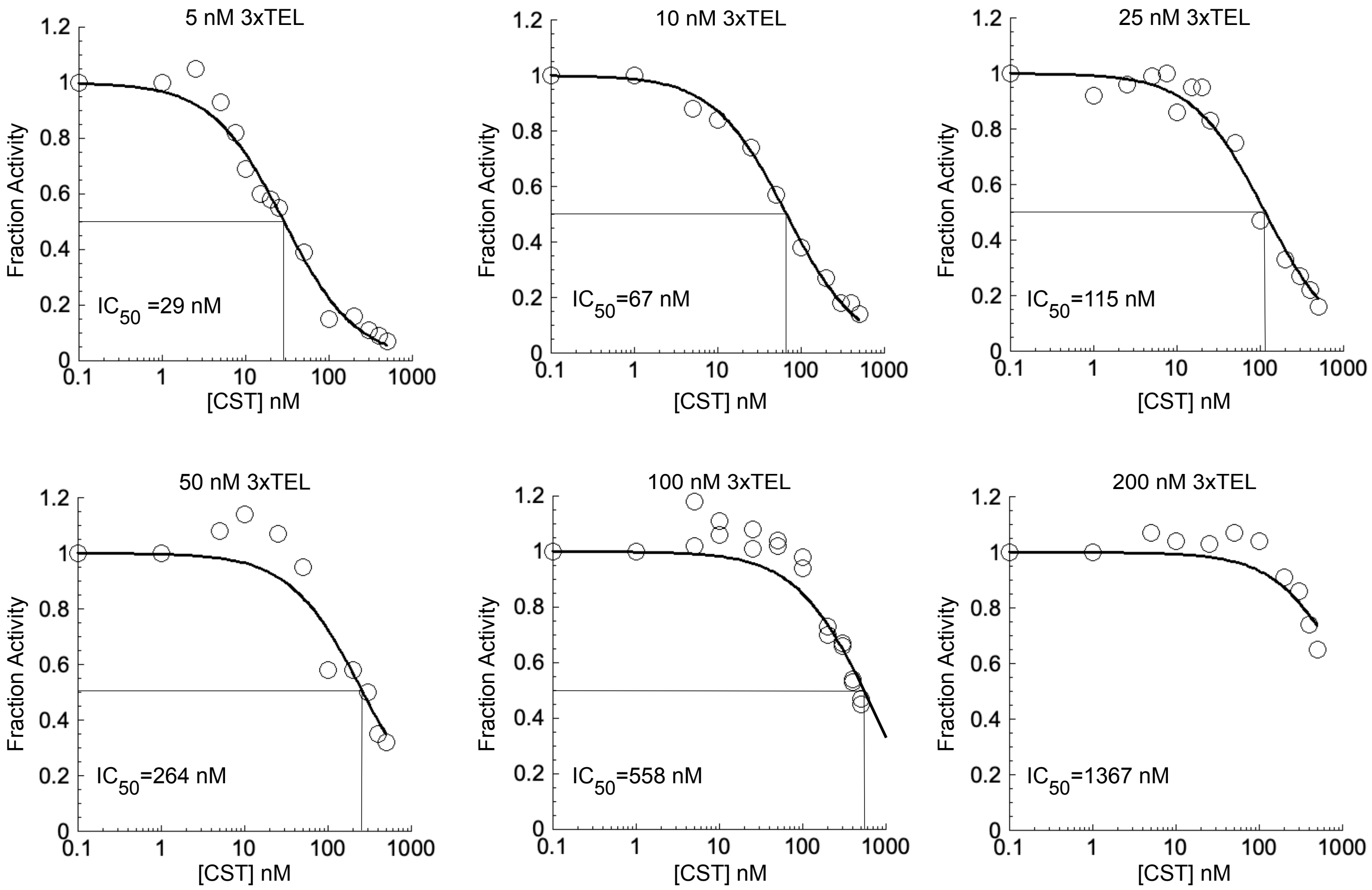


**Supplementary Figure S2**. Inhibition of telomerase activity by pre-incubation with WT CST protein. Each point represents a lane on telomerase activity gel, with total radioactivity incorporated normalized to that obtained with no CST added. 3xTEL primer concentrations varied as indicated. Data points were fit with equation 3. The 200 nM 3xTEL data do not extend to 50% inhibition, so the $IC_{50}$ is approximate.

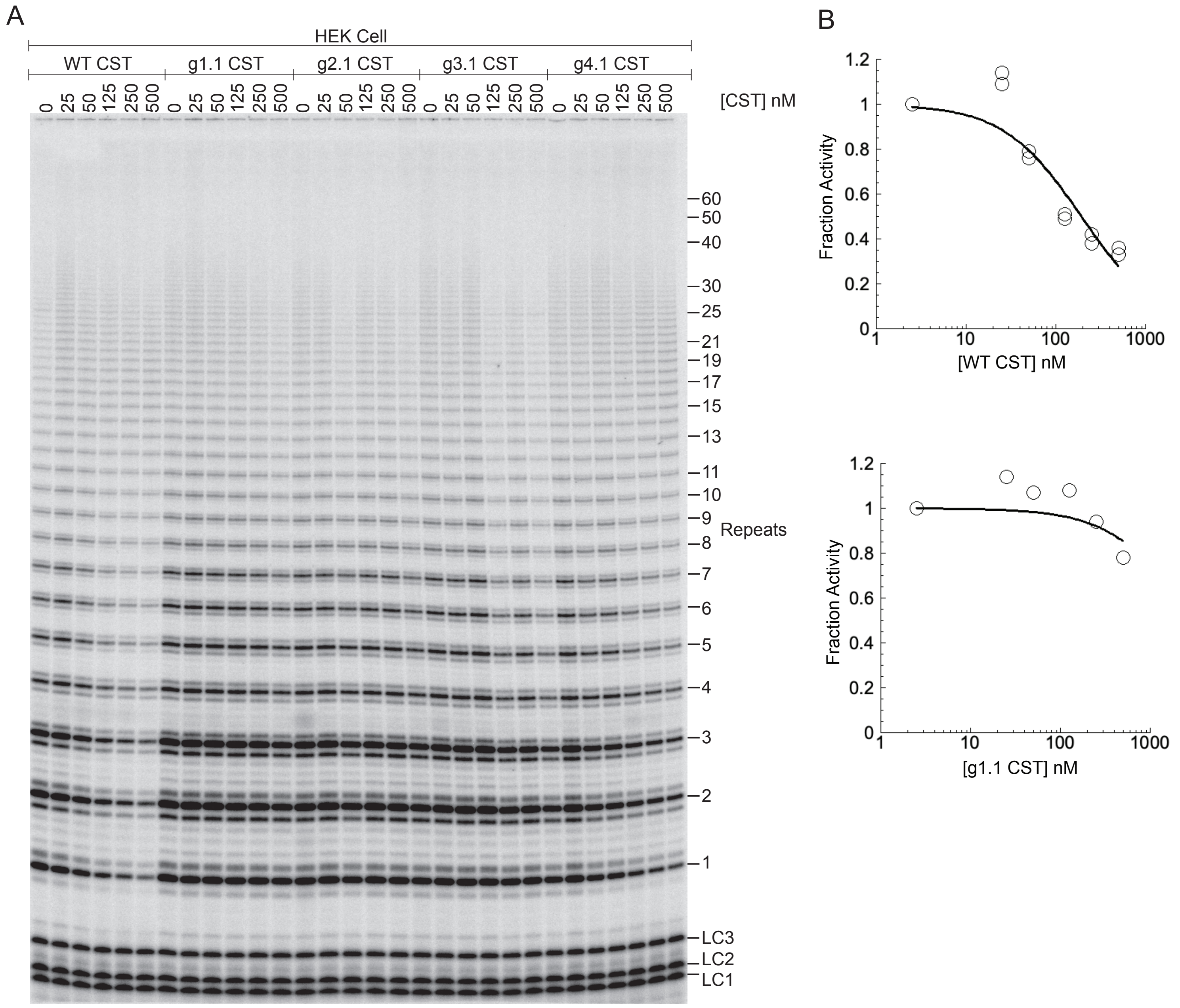


**Supplementary Figure S3**. Survey of telomerase inhibition by WT and mutant CST proteins. (**A**) Telomerase reactions with standard 10 nM 3xTEL primer for 60 min. (**B**) Quantification of WT and g1.1 inhibition, fit with equation 3; the other mutants are analyzed in more detail in Supplementary Figures S4, S5, and S6.

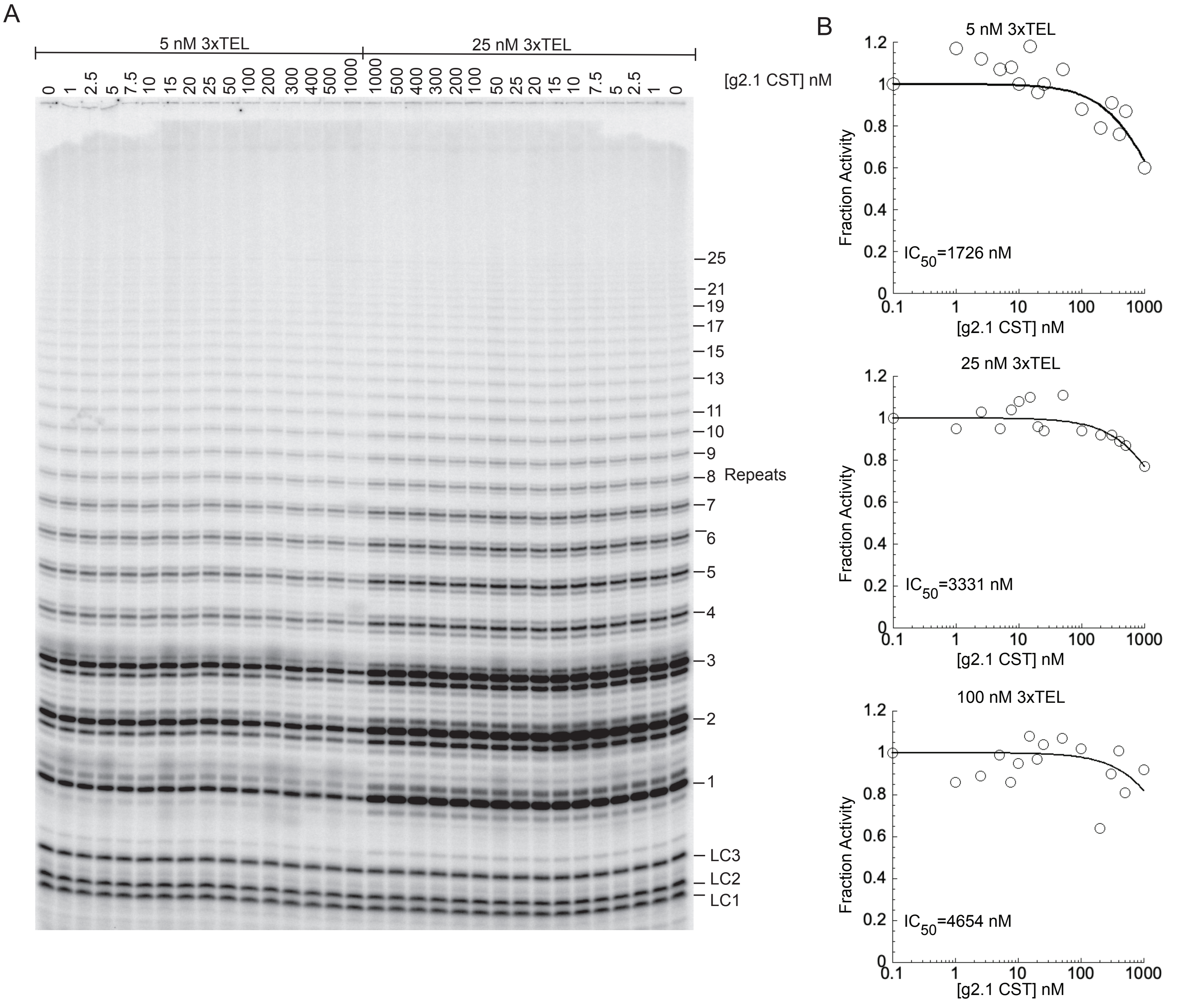


**Supplementary Figure S4**. Inhibition of telomerase activity by g2.1 mutant CST protein.
(**A**) Gel electrophoretic analysis of telomerase activity at two 3xTEL DNA primer concentrations.
(**B**) Quantification of counts in each lane of panel (A) and an additional experiment at 100 nM primer, fit with equation 3. The calculated $IC_{50}$ values at 25 and 100 nM 3xTEL are included for completeness, but they are not expected to be accurate because so little inhibition was observed at the highest CST concentrations tested.

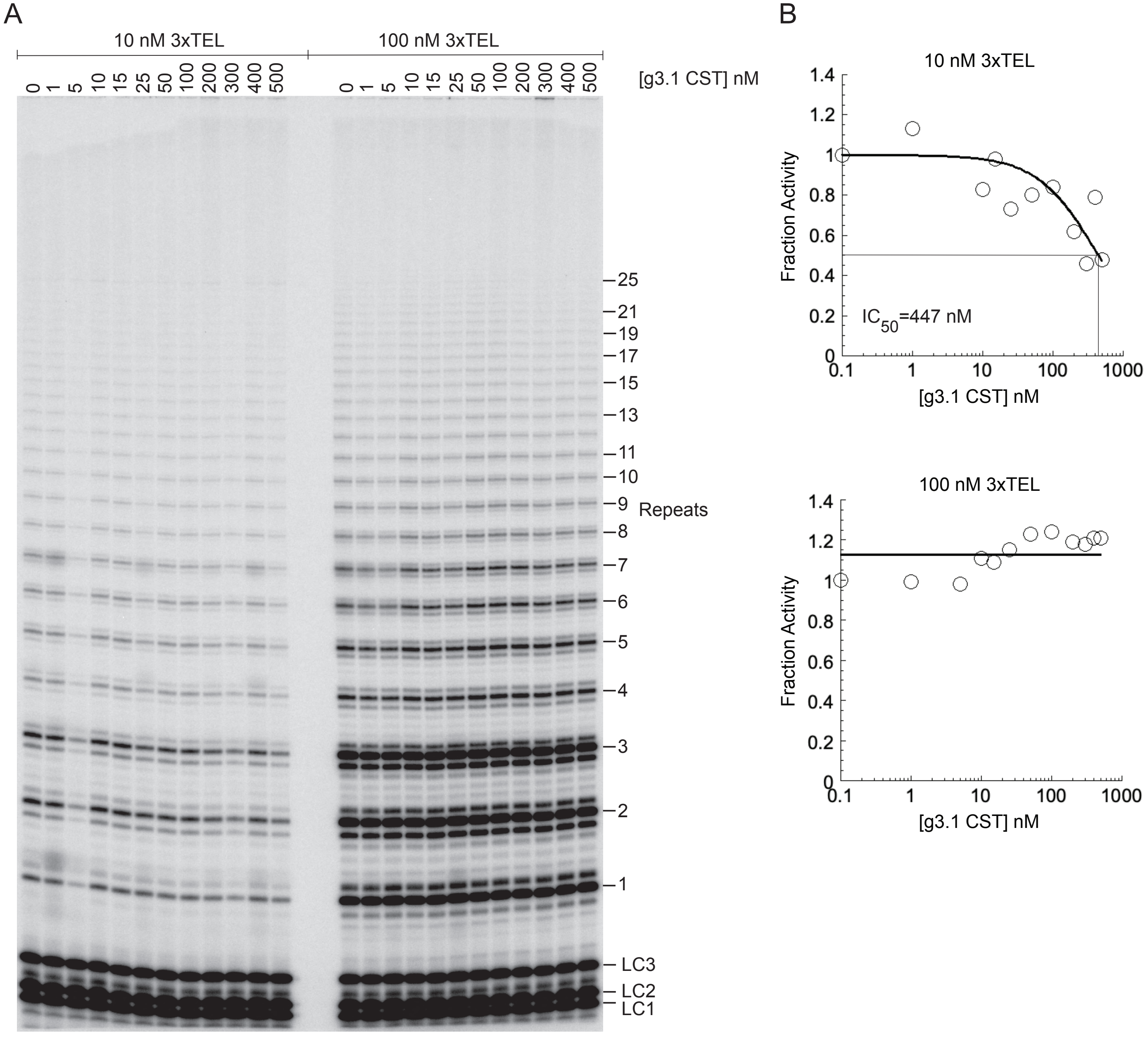


**Supplementary Figure S5**. Inhibition of telomerase activity by g3.1 mutant CST protein.
(**A**) Gel electrophoretic analysis of telomerase activity at two 3xTEL DNA primer concentrations.
(**B**) Quantification of counts in each lane of panel (A) fit with equation 3.

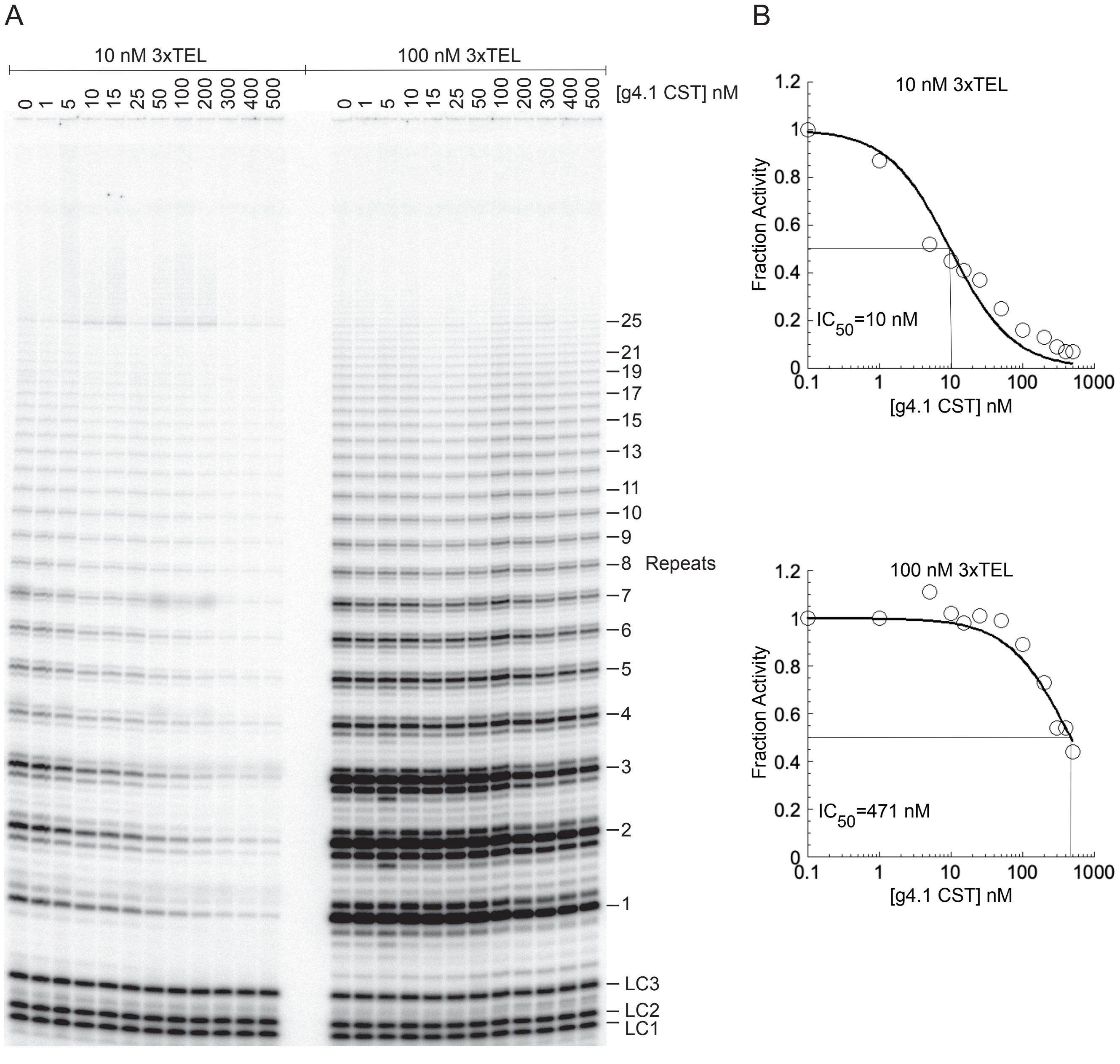


**Supplementary Figure S6**. Inhibition of telomerase activity by g4.1 mutant CST protein. (**A**) Gel electrophoretic analysis of telomerase activity at two 3xTEL DNA primer concentrations. (**B**) Quantification of counts in each lane of panel (A) fit with equation 3.

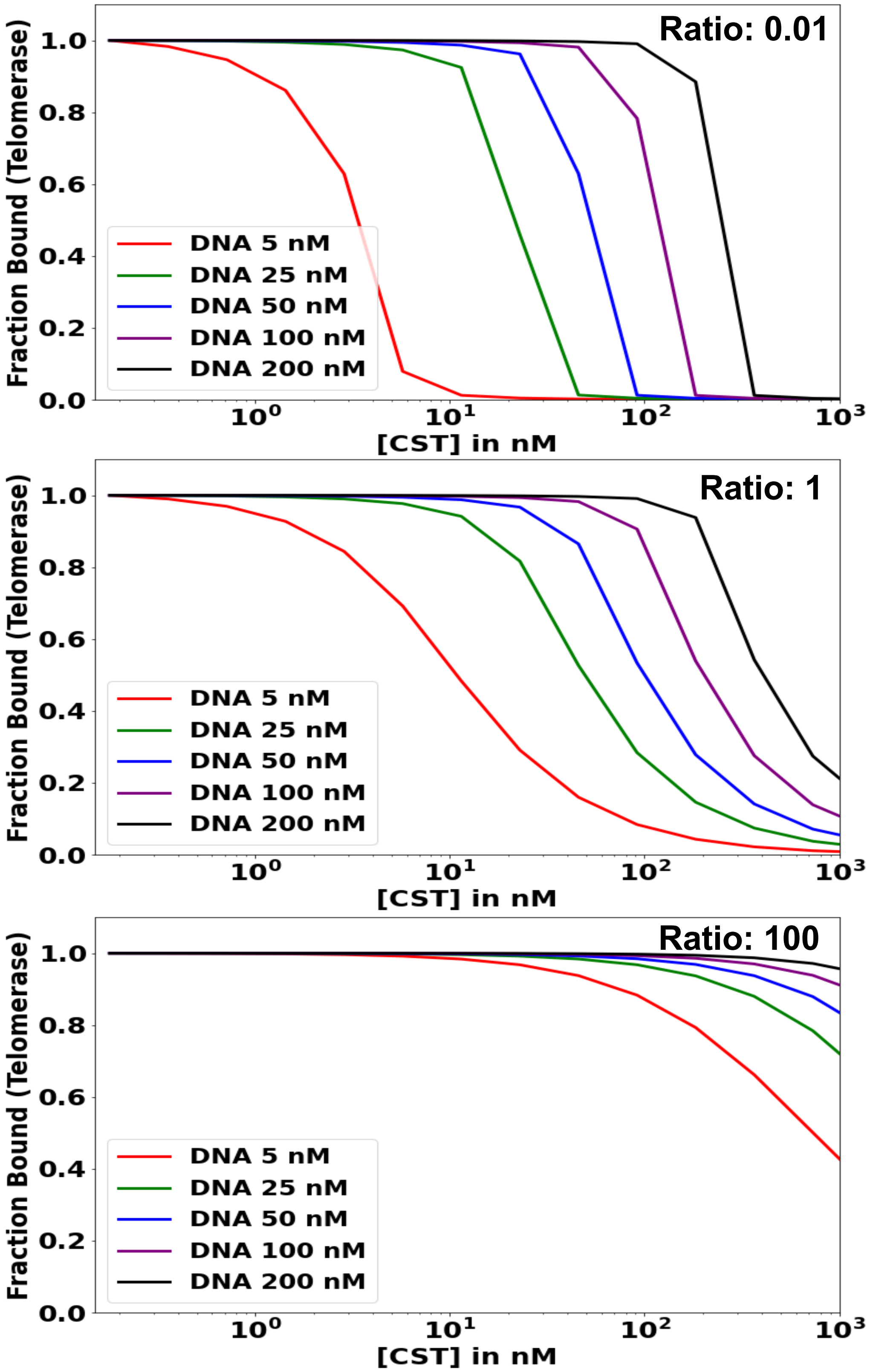


**Supplementary Figure S7:** Validation of the simulation of telomerase inhibition by CST. Simulations were performed under the following conditions: $\gamma$ = 1.0 (100% active CST), concentration of telomerase = 2.0 nM, and concentration range of WT CST from 0.0 to 1,000 nM. Predicted fraction bound curves for telomerase to DNA were plotted with initial DNA concentrations of 5.0 nM, 25 nM, 50 nM and 100 nM. This was done for three different values of the ratio of CST-DNA $K_{dB}$ to telomerase-DNA $K_{dA}$: 0.01, 1.0 and 100. The python Matplotlib graphics package (25) was used to plot the simulated data.

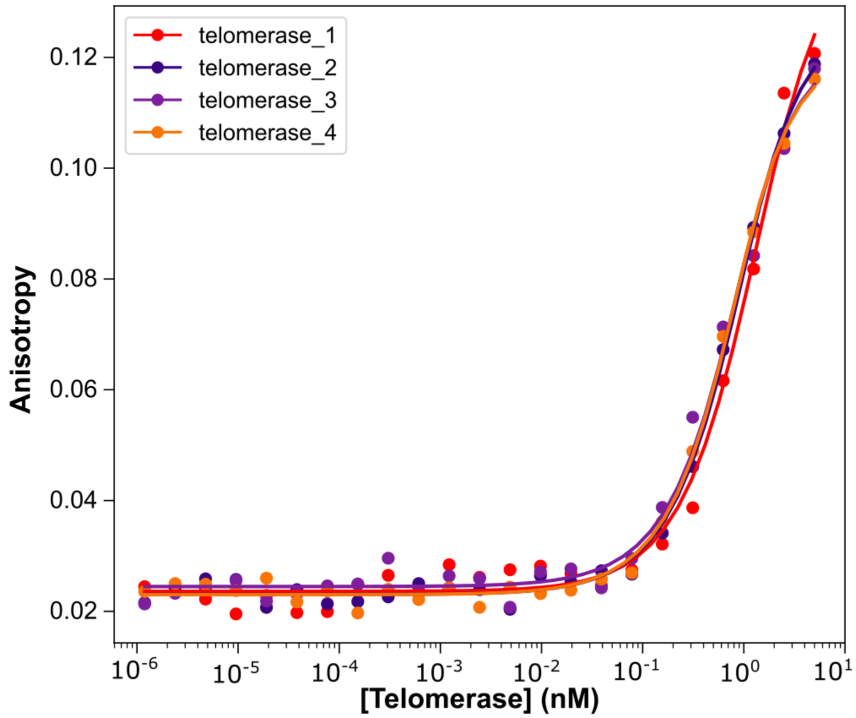


| | telomerase_1 | telomerase_2 | telomerase_3 | telomerase_4 |
|---|---|---|---|---|
| **Kd (nM)** | 0.9 | 0.48 | 0.39 | 0.37 |
| **S** | 0.121 | 0.1056 | 0.0989 | 0.0995 |
| **O** | 0.0236 | 0.0231 | 0.0245 | 0.023 |
| **R^2** | 0.988 | 0.9962 | 0.9888 | 0.9964 |

**Supplementary Figure S8.** Equilibrium binding constant of telomerase to 3xTEL DNA determined by FP.
